# Chronic Intermittent Ethanol Activates CHOP in Striatal Microglia and Exacerbates Ethanol Withdrawal

**DOI:** 10.1101/2020.09.02.280156

**Authors:** Rapheal G. Williams, Brett D. Dufour, Kevin R. Coffey, Atom J. Lesiak, William B. Nickelson, Aliyah J. Dawkins, Gwenn A. Garden, John F. Neumaier

## Abstract

Repeated cycles of alcohol intoxication and withdrawal induce profound changes in gene expression that can contribute to the physiological and behavioral consequences of ethanol. Since neuroinflammation is an important consequence of these changes, we used a novel strategy to investigate the impact of repeated cycles of chronic intermittent ethanol vapor and withdrawal on the RNAs actively undergoing translation in microglia in striatum, a key region involved in the relapse to ethanol consumption. We performed deep sequencing of the “translatome” from striatal microglia of male and female RiboTag mice, yielding a snapshot of RNA translation during alcohol intoxication and after 8 hours of withdrawal. Chronic intermittent ethanol produced robust changes in the translatome, with increases in genes and pathways associated with cytokine signaling, indicating increased neuroinflammation and microglial activation. After 8 hours of ethanol withdrawal, many inflammatory pathways remained upregulated and phagocytotic and proapoptotic pathways were increased. Using unbiased network analysis, we identified gene modules that were differentially expressed in ethanol intoxicated vs. withdrawing animals. Genes associated with the unfolded protein response (UPR) were over-represented in one such module after withdrawal, including the transcription factor Ddit3 (CHOP), an important mediator of the UPR. We tested the impact of conditional knockout of CHOP from microglia specifically; following withdrawal from chronic intermittent ethanol, these mice had reduced thermoregulatory disturbances, anxiety-like behavior, and voluntary ethanol consumption compared to wild-type littermates. We conclude that CHOP and the UPR in microglia may be important targets for reducing the impact of withdrawal from chronic ethanol exposure.

---

**C**hronic alcohol use disorder (AUD) is associated with neurotoxicity, neuroinflammation, and immune activation (Crews et al., 2017) The degree of toxicity is exacerbated both by dose and duration of intake (Harding et al., 1996; Miguel-Hidalgo et al., 2006). Furthermore, repeated attempts at alcohol cessation in dependent individuals is common before achieving sustained abstinence, and a kindling model has been proposed to explain how these repeated cycles of alcohol use and withdrawal lead to successive increases in neuroinflammation, neurotoxicity, neurodegeneration, and withdrawal symptom severity (Becker and Hale, 1993). Indeed, individuals with a history of substantial chronic alcohol use show significant levels of neuronal cell loss (Harding et al., 1996; Miguel-Hidalgo et al., 2006), astrocyte and oligodendrocyte pathology (Miguel-Hidalgo et al., 2006), microglial activation (He and Crews, 2008) and increased neuroimmune signaling (He and Crews, 2008). While there is strong evidence for how alcohol induced toxicity to neurons contributes to AUD and alcohol withdrawal symptomology (Miguel-Hidalgo et al., 2006; Badanich et al., 2011), the contributions of alcohol associated glial pathology and neuroimmune activation are less clear.

As the primary resident immune cells of the brain, microglia are key regulators in neuroinflammation. At ‘rest’, they actively surveil the brain for alterations in homeostasis detected by numerous membrane embedded receptors (Hanisch, 2002; Kettenmann et al., 2011). Thus, microglia play a dynamic and crucial role in healthy brain function, including modulating neurogenesis, neuronal circuit formation, synapse maturation during neurodevelopment, and by regulating neuronal signaling and neuroplasticity in the healthy adult brain (Salter and Stevens, 2017). However, in cases of tissue trauma, microglia will “activate,” migrating to the site of trauma, proliferating, changing their morphology (larger soma and shorter processes) and regulating the neuroinflammatory response by secreting neuroimmune and trophic factors and by phagocytosing debris (Kettenmann et al., 2011). The transition from resting to activated states is complex and involves many adaptations in microglial gene expression that may have important implications for the role of microglia in AUDs.

Alcohol use and withdrawal have profound effects of microglial function, and these alcohol-induced alterations in microglial function in turn contribute to alcohol associated behaviors and withdrawal symptomology (Crews et al., 2017). However, many questions remain regarding how microglia respond to alcohol on a molecular level, including which changes are adaptive and which exacerbate pathology. Additionally, it is unclear how these microglial alterations in turn impact alcohol induced neurotoxicity, physiological withdrawal symptoms, and behavioral consequences such as craving and impulsivity. In the series of experiments presented here, we used a bioinformatics approach to characterize the effects of alcohol intoxication and withdrawal on microglia by assessing alterations in microglia-specific gene expression. To do so, we utilized transgenic mice expressing RiboTag selectively in microglia, thus enabling specific isolation of actively translating mRNA transcripts from striatum, a method that we have used previously to identify RNAs that are co-regulated in specific cells (Levinstein et al., 2020; Lesiak et al., 2021; Coffey et al., 2022). Since Chronic Intermittent Ethanol (CIE) primes and activates microglia and induces potent neuroinflammation in exposed subjects (Crews et al., 2017), we used the well-established vapor CIE paradigm to identify the engagement of alcohol-sensitive, microglia-specific molecular pathways. The transcription factor C/EBP homologous protein (CHOP, gene symbol Ddit3) is a key activator of the unfolded protein response, a cellular stress response that contends with misfolded proteins that impair cellular health, was upregulated after withdrawal from ethanol, so we selected it for further investigation. We found that selective deletion of CHOP from microglia reduced anxiety-like behavior and voluntary ethanol consumption after CIE, suggesting that the unfolded protein response contributes to escalated ethanol consumption in this model. These results suggest that the inhibition of CHOP, and the UPR more generally, may be a valuable therapeutic approach to treating alcohol withdrawal.

## Results

### Methodological Validations

Treating Cx3cr1-Cre/Ribotag double transgenic mice (Figure 1a) with tamoxifen resulted in strong expression of hemagglutinin (HA) tagged Rpl22 in Iba1 expressing microglia (Figure 1c). Mice were then subjected to CIE vapor exposure (Figure 1b), consistently reaching blood ethanol concentrations of approximately 200 mg/dL (Figure S1). RNA extraction from one striatum per mouse yielded 468±25 ng (Mean±SEM) of RNA (this was used to prepare “Input” RNA) while immunoprecipitation of RiboTag-associated RNA from each sample yielded 59±6 ng of “IP” RNA. Input samples were pooled within group (10 ng per sample), and then 10ng of RNA from each sample was used for sequencing library preparation. RNA sequencing from IP samples yielded 6.83±0.18 million reads (Mean±SEM) while Input samples yielded 3.58±1.00 million reads. All fastq files passed FastQC basic statistics, per base quality and per sequence quality.

**Figure 1.**
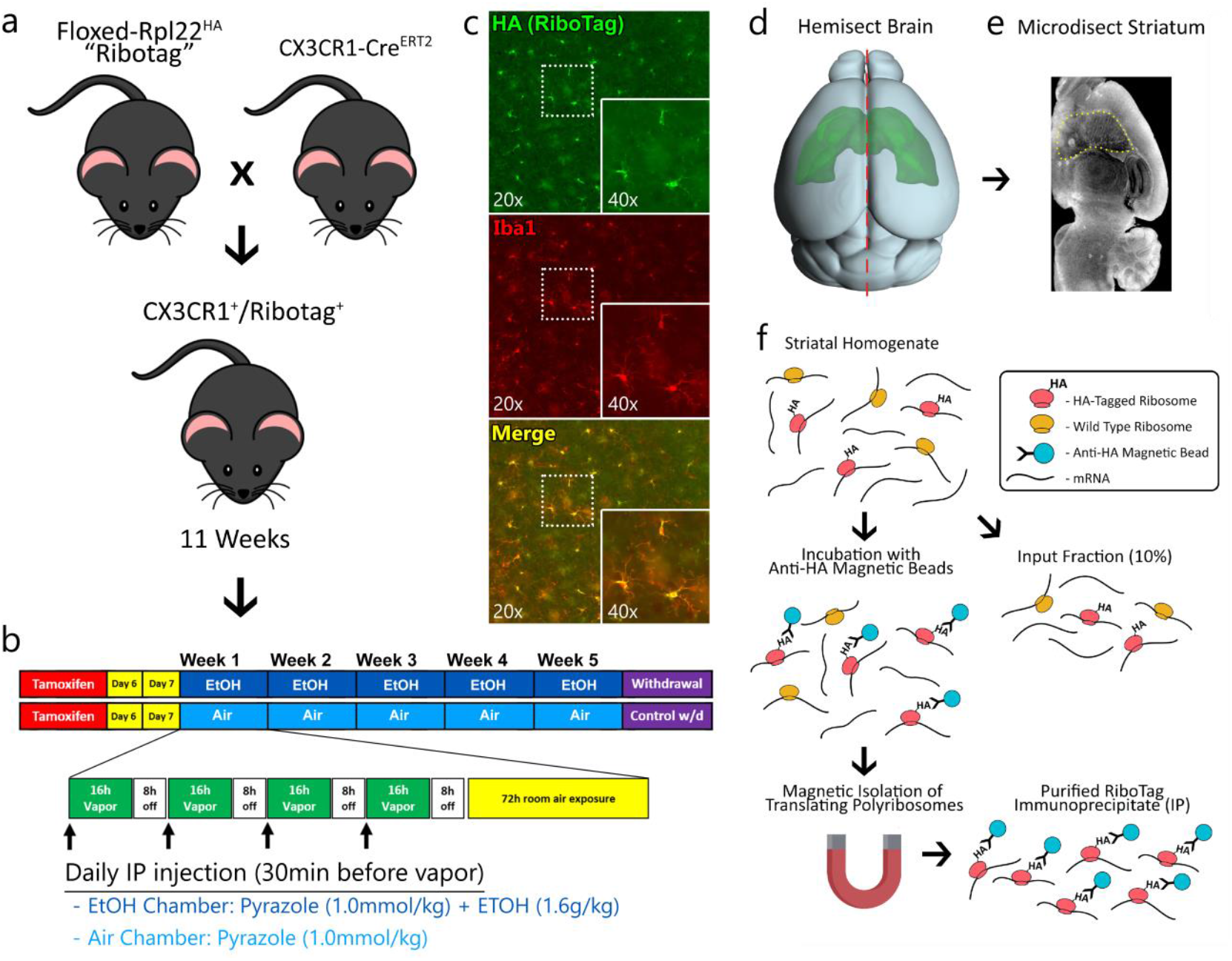
Experimental Overview. **a)** Experimental male and female mice were generated by crossing tamoxifen-inducible CX3CR1-CreERT2 hemizygous mice with homozygous floxed RiboTag mice. **b)** The experiment was performed as a 2×2 treatment design (n=6/group) with all mice receiving tamoxifen for 5 days at 14 weeks. At ∼15 weeks of age the mice underwent chronic intermittent ethanol exposure (or air exposure). Animals were sacrificed after the week-5 session at either 0 hours or 8 hours after the final exposure. **c)** Brains from these animals express HA-tagged ribosomes exclusively in microglia. **d**,**e**) The brains were rapidly extracted, hemisected, and the striatum was microdissected and homogenized. **f)** RiboTag isolation procedure involves incubating tissue samples with anti-hemagglutinin antibody magnetic beads and performing magnetic isolation of RiboTag-positive, ribosome bound RNAs. Scale bar = 20 µm.

### Microglia Specific Markers are Highly Enriched in RiboTag-Seq Samples

Differential expression analysis of IP vs Input samples isolated 2,418 genes significantly enriched in the IP samples. Among the highly enriched genes are numerous classic microglia markers such as Itgam (cd11b), Csf1r, Mpeg1, and CX3CR1 (Figure 2a). Gene set enrichment analysis (GSEA) of IP vs Input samples further validated our enrichment of classic microglia pathways, such as cytokine binding, purinergic receptor activity, and NF-kB binding (Figure 2b). This procedure is consistent across experiments; 80% of the differentially expresed genes in the present study were also differentially expressed in our previous CX3CR1-Ribotag study (Coffey et al., 2022) (Figure 2c). Immunoprecipitation and sequencing protocols were identical across these two studies, but animals were exposed to morphine instead of ethanol. Despite this difference, IP samples show highly similar enrichment of microglia specific genes. Further, genes that were highly enriched (log2(fc) > 4) shared nearly 100% concordance across experiments (Figure 2d).

**Figure 2.**
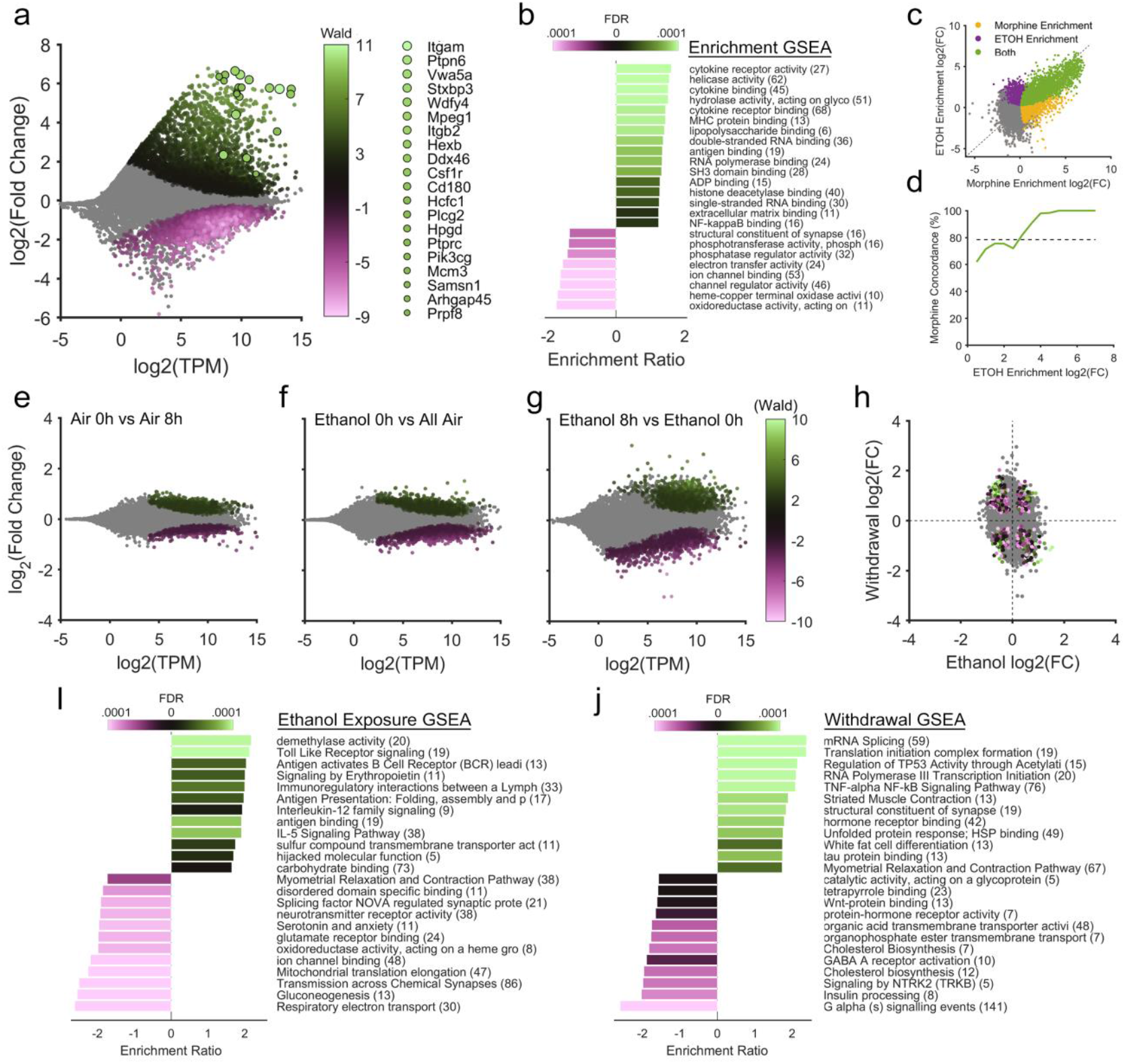
Differential Expression Analysis. **a)** The RiboTag-Seq samples were dramatically enriched with microglial markers such as Itgam (cd11b) and CX3CR1. Color represents enrichment statistic (Wald) for genes with an FDR< 0.1) while circle size represents relative significance value (FDR). **b**) GSEA reveals an upregulation of genes associated with classical microglial gene sets such as “cytokine binding” and “purinergic receptor activity”. **c**) 80% of the differentially enriched genes in the present study were differentially expressed in our previous CX3CR1-Ribotag study assessing microglial changes in gene expression during morphine tolerance and withdrawal 31. **d**) Genes that were highly enriched (log2(fc) > 4) shared nearly 100% concordance with our previous CX3CR1-Ribotag RNA-Seq Study. **e**) Extraction time alone produced relatively few differentially expressed genes in air exposed controls. **f**) Ethanol exposure produced a modest number of upregulated and downregulated DEGs as compared to air exposed animals. **g**) Ethanol withdrawal produced dramatic differential gene expression in striatal microglia. **h**) Gene expression changes during withdrawal were not correlated to gene expression changes during ethanol exposure. GSEA was performed using several databases, including GO: Molecular Function, Reactome, and Wiki Pathways. The top 4 upregulated and downregulated gene sets from each database are displayed. **i)** Ethanol exposure upregulated classic microglial activation gene sets such as TLR signaling, IL12, and IL5 signaling, **j**) while ethanol withdrawal upregulated gene sets related to neuroinflammation such as TNF-alpha/NF-kB signaling, and gene sets associated with cellular stress such as the unfolded protein response and heat shock proteins.

### Ethanol Withdrawal Produce Dramatic Differential Gene Expression in Microglia

There were few differentially expressed genes (DEGs) in microglia between the two air exposure control groups (Air-0h vs Air-8h; Figure 2e). Ethanol exposure produced a modest 151 upregulated and 288 downregulated DEGs (Figure 2f). By contrast, withdrawal produced nearly double the number of DEGs, with 468 upregulated and 354 downregulated (Figure 2g). There was no consistent relationship in the direction or magnitude of change in DEGs when comparing ethanol exposure (EtOH-0h vs all Air) to withdrawal (EtOH-8h vs. EtOH-0h, Figure 2h).

### Gene Set Enrichment Analysis of Microglia RNA During Ethanol Exposure and Withdrawal

GSEA provides many insights into how typical microglial function (i.e. phagocytosis, inflammatory signaling, etc.) was altered by ethanol and withdrawal. Relative to air treated controls, ethanol exposure (EtOH-0h) resulted primarily in an upregulation in cytokine signaling related gene sets including Toll-Like Receptor (TLR), Interleukin-5 (IL-5), and Interleukin-12 (IL-12) family signaling sets, all indicative of increased neuroinflammation and microglial activation (Figure 2i). Type II interferon (INF) signaling was also regulated; and much like TLR4 pathways, IFN cascades activate a broad neuroinflammatory response (Monteiro et al., 2017). The Microglial Pathogen Phagocytosis Network and the TYROBP network (Konishi and Kiyama, 2018), both associated with active phagocytosis, were also upregulated by ethanol exposure. The Apoptosis gene set was also upregulated, suggesting that there was an increase in microglial programmed cell death. Ethanol exposure was also associated with a downregulation in cellular metabolism gene sets, including Glycolysis and Gluconeogenesis (Figure 2i), TCA Cycle, Amino Acid Metabolism, Oxidative Phosphorylation, and Electron Transport Chain sets. While this likely reflects ethanol’s well-known interference with metabolic pathways (Manzo-Avalos and Saavedra-Molina, 2010), it also suggests that ethanol induced metabolic impairment may negatively impact the functional integrity of microglia. Relative to the acute ethanol intoxication timepoint (EtOH-0h), numerous gene sets were exclusively altered during withdrawal (EtOH-8h, Figure 2j). There was an upregulation in the pro-inflammatory TNFα/NFkB cytokine gene set, as well as an upregulation in cellular stress related gene sets such as Heat Shock Protein binding and the UPR. Finally, numerous G-protein coupled receptor gene sets were downregulated during withdrawal. Only the top gene 12 upregulated and downregulated gene sets following affinity propagation redundancy reduction are displayed in Figure 2. The compete Ethanol GSEA analysis can be viewed interactively in our data repository.

### WGCNA Reveals the Induction of the Unfolded Protein Response in Striatal Microglia during Ethanol Withdrawal

A minimum spanning tree of the entire RiboTag-Seq matrix and corresponding module labels were projected into 2D space using a force directed layout (Figure 3a). The statistical significance of increased and decreased DEGs in EtOH-8h vs EtOH-0h is overlaid on the minimum spanning tree to visualize the relationship between modules and individual gene significance (Figure 3b). Several modules of genes were upregulated in EtOH-8h animals (Figure 3c), suggesting that the genes in each module may have been upregulated in a coordinated fashion. Even though the clustering is not guided by any a-priori considerations about gene function, the fact that highly significant DEGs fell into only four gene modules in the EtOH-8h vs. EtOH-0h conditions suggests that these modules provide useful clues in identifying gene modules that were perturbed by ethanol withdrawal.

**Figure 3.**
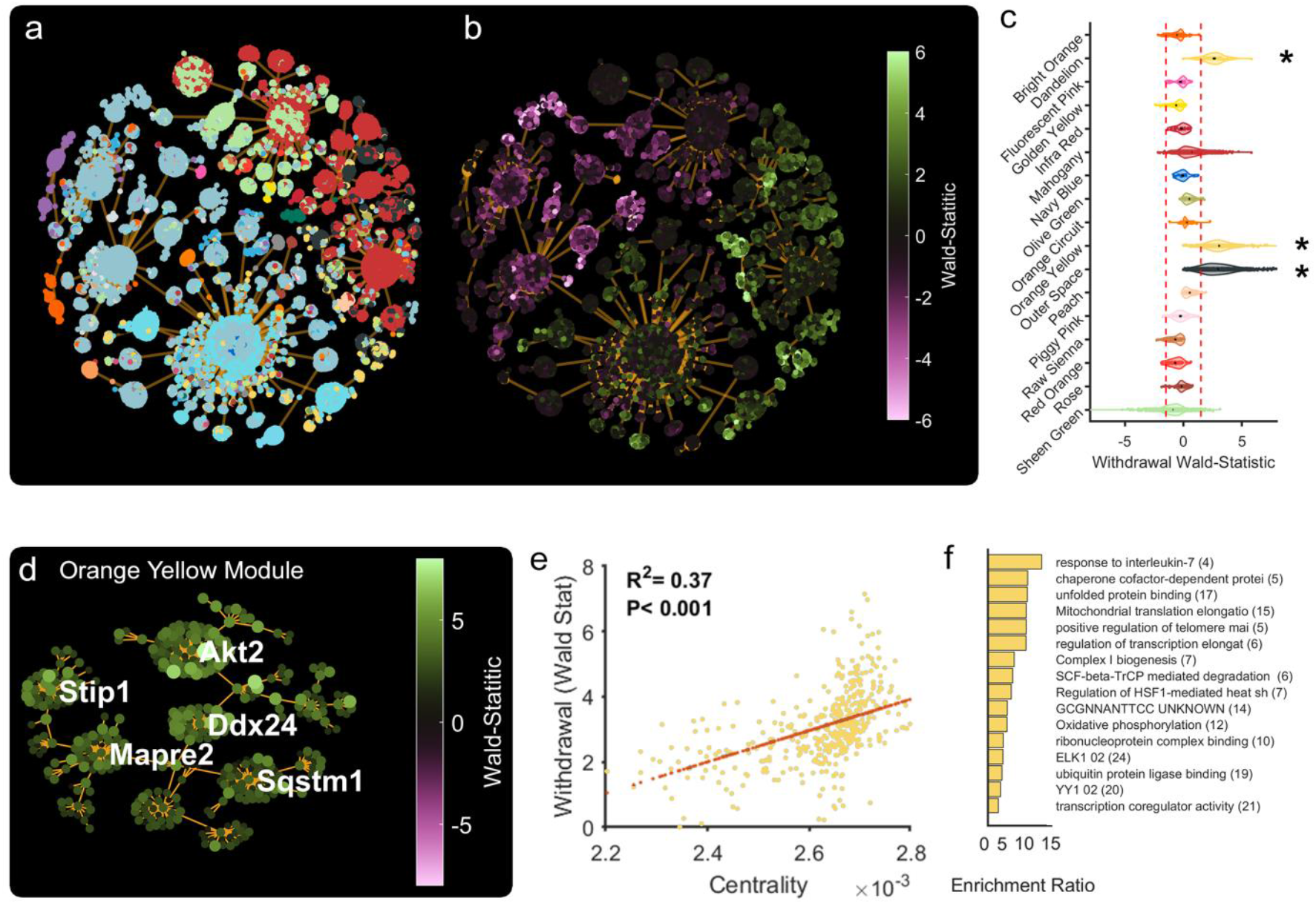
WGCNA Analysis of Ethanol Withdrawal. **a**) Genes are coded by module color and projected into a 2D minimum spanning tree using a “force” layout. **b**) The same genes are coded by differential expression (Wald-statistic) data during ethanol Withdrawal. **c**) A subset of WGCNA modules are plotted based on their differential expression metrics (Wald-statistic). Modules who’s mean Wald-statistic was less than -1.5 or greater than 1.5 were flagged for further analysis here or in Figure S2. **d**) The Orange-Yellow module projected into 2D space using a minimum spanning tree and showing the top five most central genes. **e**) Centrality (a measure of network importance) in the Orange-Yellow module is correlated to differential expression for withdrawal. **f**) Over-representation analysis on the Orange-Yellow module shows that genes related to IL-7 response, the unfolded protein response (UPR) and others are upregulated during ethanol withdrawal in striatal microglia.

For example, the “Orange-Yellow” was examined to determine if network membership conferred functional relevance to withdrawal related differential expression. An undirected network graph was generated for the Orange-Yellow module and was projected onto 2D space using a minimum spanning tree (Figure 3d). Each gene’s importance within the network is defined using a centrality metric, calculated using “closeness” for the distance calculation, and the sum of each node’s edge weight (similarity) as the “cost” function. For the Orange-Yellow module, network centrality was highly correlated to individual gene differential expression during withdrawal (Figure 3e). Over-representation analysis (ORA) was used to determine if the genes in the Orange-Yellow module likely originated from any known gene pathways (Figure 3f). ORA was performed using WebGestalt (Liao et al., 2019) to search Biological Process, Molecular

Function, Kyoto Encyclopedia of Genes and Genomes (KEGG), Reactome, and Transcription Factor Target gene sets. One substantial and significantly overrepresented pathway in the Orange-Yellow network involves genes engaged in the UPR. This is consistent with the Molecular Pathway GSEA results that indicated an upregulation in the UPR during withdrawal (Figure 2i,j). The UPR is activated in response to an accumulation of unfolded or misfolded proteins in the lumen of the endoplasmic reticulum (ER) and is a highly conserved cellular stress response related to ER stress. The UPR can be a homeostatic response to return the system to baseline but may also lead to stress-induced apoptotic cell death. There are three major ER sensors that can initiate this system: PERK (Eif⍰k3), ATF6, and IER1 (Ern1). IER1 was significantly increased in EtOH-0h mice compared to Air controls and is associated with early time points of UPR activation, whereas ATF4 is considered a later phase of activation (Nishitoh, 2012) and was increased in EtOh-8h animals during withdrawal. Since ATF4 is a key activator of the transcription factor C/EBP homologous protein (CHOP, gene symbol Ddit3), we examined the level of ribosome associated Ddit3 mRNA and found that it was significantly increased by withdrawal (q=0.002). Other gene modules may also be of interest but were not analyzed deeply in this report and can be reviewed in the supplementary figures and our code repository (Figure S2).

We also examined microglial morphology, and several mRNAs and proteins localized in microglia, using histological approaches in wildtype C57Bl/6J mice following CIE exposure and withdrawal. We focused on the same 0 and 8 h timepoints that were used for RNA analyses. Thus, C57Bl/6J mice received ethanol or air exposure and were sacrificed at 0 or 8 hrs after completing the final CIE session. We performed fluorescent in-situ hybridization, RNAscope, followed by IHC to localize mRNA of CHOP (Ddit3) and Dnaja1, which codes for DNAJA1, a protein that serves as a chaperone for heat-shock proteins (i.e. HSP70) and is also involved in the UPR. We first measured several morphological features in Iba-1 immunostained microglia for each treatment group. There were no differences between groups in the general striatal microglial morphology, including no changes in microglia number, surface volume, process branch points or complexity (Figure S3c-g). RNAscope quantitation of CHOP and Dnaja1 in situ hybridization signal localized to microglia did not differ between groups, regardless of ethanol exposure (Figure S4a-h). To further characterize the effects of ethanol withdrawal on protein expression of major molecular substrates of the unfolded protein response (UPR), and proteins related to the results seen in our RNA sequencing data, immunohistochemistry was performed on striatal tissue extracted from C57Bl/6J mice after 5 weeks of the CIE paradigm immediately following ethanol exposure and 8 hours into withdrawal. We measured signal intensity within microglial cells after staining for CHOP, grp78, IRE1α, Akt2, and Stip1 immunoreactivity although none of these effects were significantly different at the 8-hour timepoint (Figure S4i-n). Finally, since the time course of transcriptome, translatome, and protein levels can differ, we used RTqPCR to validate some of the RNA findings in a new cohort of Cx3cr1-Cre/Ribotag double transgenic mice (n=12) exposed to CIE to measure several of the RNA targets in the microglia-specific IP samples as well as the total RNA from striatum (INPUT) immediately (0 hours), 8 hours, and 24 hours following their final exposure on week 5. We investigated three genes of interest were identified by the RNA sequencing data: CHOP, Akt2, and Stip1. For CHOP, there was an overall effect of RNA source, and a time by source interaction (F=5.64, p=0.026, GLMM), with the microglia-derived (IP) CHOP RNA being differentially expressed across treatment conditions (Figure 4a) with post-hoc analysis (Dunnett’s test) showing that CHOP RNA was increased at 24 hours of withdrawal (t=3.23, p=0.035) while there was an increase trending towards significance at 8 hrs (t=2.85, p=0.064). Input RNA from the same subjects did not reveal a difference in CHOP across withdrawal timepoints, suggesting the effect of ethanol on CHOP was microglia specific. The differences across time points for both IP and input RNA for Akt2 and Stip1 were not significant (Figures 4b,c).

**Figure 4.**
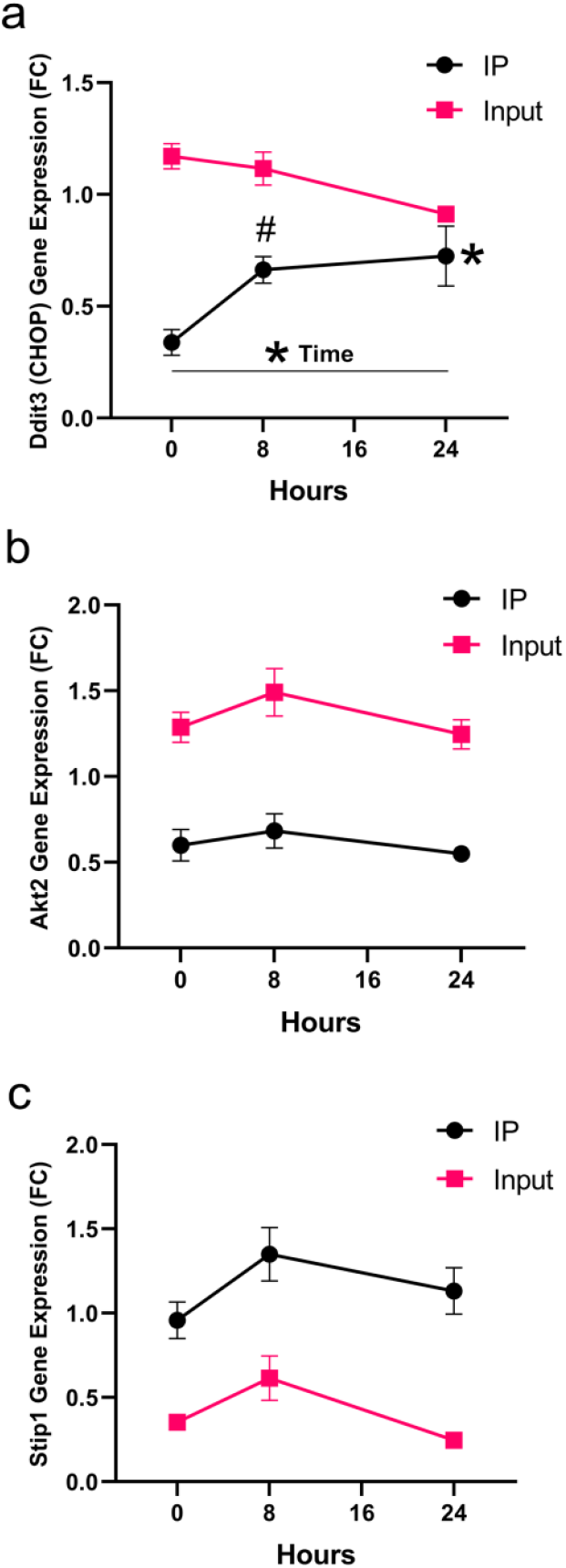
CHOP mRNA actively undergoing translation increases across early alcohol withdrawal. RNA was purified from striatum from mice expressing RiboTag in microglia for both microglial translatome (IP, black symbols) and total striatal RNA (Input, pink symbols) at 0, 8, and 24 hours after completion of final CIE session. **a)** CHOP (Ddit3) RNA, **b)** akt2 RNA, and Stip1 RNA. * p<0.05, # p=.064 trend. F values and precise post-hoc p values are in the Results section.

### CHOP Deletion in Microglia Influences Thermal Regulation, Behavior, and Voluntary Drinking Phenotypes

Since the RNAseq results implicated the UPR and we found evidence of altered CHOP expression during withdrawal, we probed this system’s involvement in microglia during chronic ethanol exposure by conditional knockout of CHOP from microglia. Cx3cr1-Cre/CHOP double transgenic and Cre-negative littermate controls were utilized to analyze the effect of CHOP conditional knockout in microglia on ethanol withdrawal-related behaviors following CIE exposure. Animals were split, counterbalanced by initial body weight into either receiving air (n=10) or ethanol (n=13) for the 5 weeks of CIE exposure. These two groups were then split by genotype to analyze the effects of conditionally knocking out CHOP in microglia. Body weights were taken three times a week to ensure subject health. Weights in all groups increased in a typical manner each week as evidenced by a significant increase in weights for all groups [F(2,36)=8.96, p<0.0001, GLMM, Figure 5a] but there were no significant differences between genotypes or treatments. Hyperthermia can occur during ethanol withdrawal, so rectal body temperature was taken three times a week: once before the CIE exposure, again immediately following the final CIE exposure for the week, and then 7-8 hours after CIE exposure. Measurements were taken every week, but only the first and last weeks’ data is shown.

**Figure 5.**
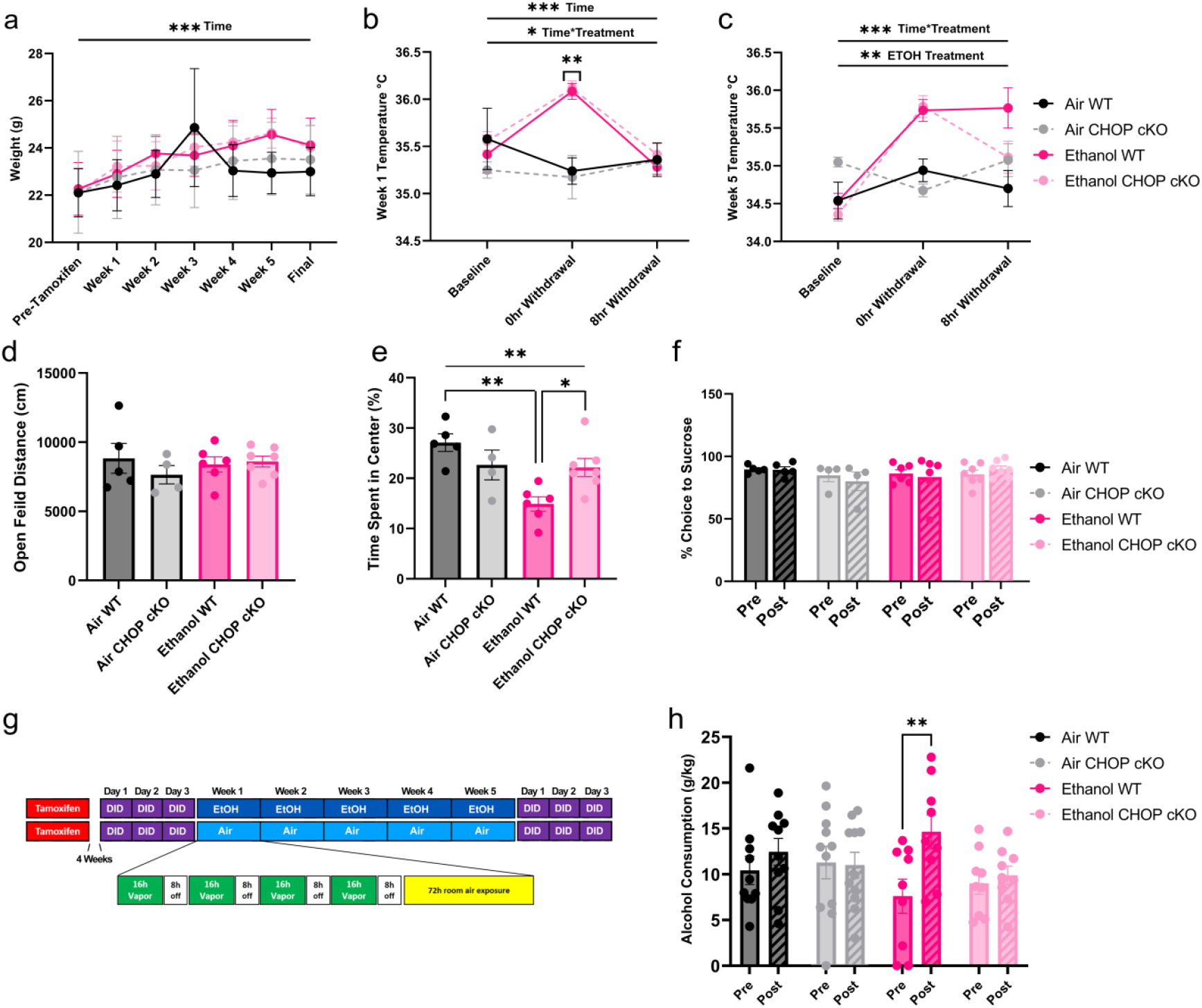
Conditional knockout of CHOP from microglia blunts withdrawal. CHOP conditional knockout (cKO) was achieved by crossing Cx3cr1-Cre and floxed CHOP mice; resulting double transgenic mice were compared to their littermate controls that did not express Cre after exposure to CIE or air control as described in the Methods section. **a)** Body weight gain was similar across all treatment groups. **b)** During the first week of CIE, ethanol-vapor treated mice were hyperthermic at the end of the ethanol vapor session, but their temperatures returned to normal at 8 hours of withdrawal; both wild-type and CHOP cKO mice responded similarly. c) At the end of the fifth week of CIE, hyperthermia was sustained after 8 hrs of withdrawal in wild-type mice but partially normalized in CHOP cKO mice. **d)** In a different cohort of mice, locomotor behavior in the open field test 48 hrs after the final vapor exposure session was similar across all treatment groups, **e)** but center time was decreased in wild-type mice that experienced CIE whereas CHOP cKO mice were not different than air-treated control mice of either genotype, indicating that the wild-type CIE mice had greater anxiety-like behavior during withdrawal. **f)** There were no differences in sucrose preference between treatment groups. **g)** Design of Drinking in the Dark experiment. **h)** Wild-type mice increased their alcohol after undergoing CIE but CHOP cKO mice did not increase voluntary ethanol consumption after CIE. *p<0.05, **p<0.01, ***p<0.001; main effects and interactions are indicated in each panel accordingly and Tukey’s post-hoc tests are indicated between groups and F values and precise post-hoc p values are in the Results section.

Thermoregulation was impacted by ethanol exposure during the first week as evidenced by an effect of time [F(2,36)=17.2, p<0.001, GLMM] and a time by treatment interaction [F(2,36)= p=0.0277, GLMM]. Post hoc analysis revealed that these changes in thermoregulation were caused by a robust increase in temperature for the animals in the ethanol groups at the 0h withdrawal time point for both genotypes (Figure 5b). By week 5, the hyperthermia from ethanol was sustained after 8 hours of withdrawal in wild-type mice, but reduced in CHOP cKO mice, with treatment by time interaction [F(2,36)=15.34, p<0.0001, GLMM], and an EtOH treatment effect [F(1,18)=10.12, p=0.0052, GLMM].

CHOP cKO blunted the prolonged hyperthermic effect as evidenced by a trend for decrease in rectal temperature at 8 hrs after the CIE exposure (p=0.0949, GLMM; Figure 5c).

We next explored whether microglial CHOP cKO impacted behavioral phenotypes of withdrawal, anxiety, and anhedonia following the final day of CIE. Forty-eight hours following their alcohol exposure behavior was observed in the open field test. Total locomotion, measured as distance traveled, did not differ between exposure groups or genotype during the 30-minute session (Figure 5d). Time spent in the center was reduced by ethanol exposure [F(3,18)=7.00, p=0.0026, GLM] and Tukey’s post hoc analysis revealed that ethanol-exposed wildtype animals spent significantly less time in the center compared to both CHOP-cKO animals (p=0.0439) and their air-exposed counterparts (p=0.0015; Figure 5e), which is consistent with withdrawal-induced anxiety-like behavior. In contrast to the effects seen in the open field test, our sucrose preference assays were not impacted by either ethanol exposure or genotype (Figure 5f).

We then used the drinking in the dark (DID) procedure (Figure 5g) to investigate the effect of conditional knockout of CHOP from microglia on voluntary ethanol consumption. There was no difference in baseline ethanol consumption between groups prior to CIE, but there was an overall effect of time [F(1,35)=5.36, p=0.0263, GLMM] between the pre-test and post-test; i.e. before and after the CIE or air exposure (Figure 5h), and a post hoc analysis confirmed that wild-type mice significantly increased ethanol consumption from pretest to posttest sessions only after CIE (p=0.0253) while microglia CHOP cKO mice did not increase consumption after either EtOH vapor or air-control treatments (Figure 5h). In summary, we found that, following withdrawal from CIE, conditional knockout of CHOP from microglia ameliorated the anxiety-like behavior, hyperthermia, and escalated voluntary ethanol consumption.

## Discussion

These studies demonstrated that there were dramatic changes in the microglial translatome — RNAs actively undergoing translation — in striatal microglia of mice that were exposed to repeated cycles of ethanol vapor exposure and withdrawal. Using deep sequencing of RiboTag-enriched RNA from microglia, we used several strategies to analyze the effect of chronic ethanol exposure and withdrawal on differential expression within the microglial translatome and found novel changes including an increase in RNAs associated with the UPR. Furthermore, we found that the conditional knockout of CHOP, an important component of the UPR, mitigated the impacts of CIE on withdrawal and voluntary ethanol consumption.

The RiboTag procedure yielded a dramatic enrichment of genes associated with microglia as compared to total RNA in the striatum (Figure 2a-d). One advantage of the method used here is that the RiboTag procedure provides a snapshot of RNA actively undergoing translation at the time of brain extraction and is derived from all ribosomes in the cell type expressing the RiboTag protein. Thus, there is no loss of RNA due to cell fractionation and sorting, preserving RNA from ribosomes throughout the cell cytoplasm and no change in RNA expression due to altered gene expression induced by the cell sorting procedure. Furthermore, we reasoned that RNAs that were translated together might also be co-regulated by the experimental manipulations, providing new insights into functional networks of genes within microglia during ethanol dependence and withdrawal. Accordingly, while our results are consistent with other studies investigating glial-associated CIE induced alterations in gene expression using alternative methods (Osterndorff-Kahanek et al., 2015; Erickson et al., 2019), we believe that RiboTag enabled us to extend the findings of these previous studies by identifying additional novel alcohol sensitive genes networks.

Using DESeq2, we found that there were many changes in individual gene expression levels between ethanol treated animals and air controls, and between ethanol exposed animals at 0 hours (intoxicated) vs. 8 hours (withdrawal) after discontinuation of ethanol vapor exposure. Hundreds of genes were either up or downregulated between treatment conditions (Figure 2e-h). These individual changes provide a great deal of data and potential leads for subsequent analysis and are presented in the supplemental data.

We next used GSEA to produce a more integrative analysis of gene changes. GSEA is a method that uses previously curated sets of functionally defined genes and examines whether there is a statistical over-representation of genes associated with these gene sets in any of the experimentally manipulated treatment groups. We found a pattern of changes that suggested the induction of inflammatory and cytokine signaling in microglia after repeated cycles of ethanol exposure (EtOH-0h), as well as an upregulation in the TNF cytokine gene set exclusively at 8 hours of withdrawal from ethanol vapor, a timepoint that has previously been associated with intense biological and behavioral signs of ethanol withdrawal (Metten et al., 2018). Although GSEA is a very sensitive strategy for detecting meaningful patterns of gene changes, it is based on the observations that were used to curate the gene sets in the first place. Therefore, we also used WGCNA, an unbiased clustering strategy that detects the relatedness amongst all the genes measured across all the individuals sampled in the study. This approach complements the other two strategies but may be useful in identifying novel patterns of gene changes and potentially novel genes that have not been functionally defined yet. WGCNA of this dataset revealed robust modules of genes which had similar expression patterns and there may be many insights to be gleaned from examining several of these modules. The “Orange-Yellow” module was particularly intriguing as it was differentially regulated in ethanol intoxication vs. withdrawal. Using the DEG analysis generated by DESeq2, we found that the Wald score for each gene in this module was strongly correlated with its centrality in the Orange-Yellow module, suggesting that this module was meaningfully associated with key biological processes that occurred in response to ethanol withdrawal. It is possible that both mediators and mitigators of withdrawal were identified. The full list of genes in this module reflect a range of genes that encode both well-known and novel proteins and the most central genes of this module reflect the coregulation of the entire module. These most central genes have been framed as “hub” genes in other studies (Lorsch et al., 2019) although they may or may not directly regulate the other genes with which they are correlated. To further interpret this gene module, we performed over-representation analysis on this gene set (Figure 3f), which is analogous to GSEA, and we found there were gene sets relating to the UPR. This result is intriguing because the UPR has been associated with ethanol toxicity and withdrawal in other tissues (Kaplowitz and Ji, 2006) and other cell types including macrophages (Kaphalia et al., 2019) but has not been detected in microglia in the central nervous system during ethanol withdrawal. Interestingly, Erickson and colleagues detected altered expression of a module involving UPR in both cortical microglia and astrocytes in acutely intoxicated mice after CIE whereas we observed this module to be reduced in EtOH-0h animals compared to air controls but increased after withdrawal (Erickson et al., 2019).

The UPR is a cellular response to stress that can be initiated when there is increased detection of misfolded proteins in the endoplasmic reticulum or mitochondria. The UPR can be a homeostatic response to return the system to baseline but may also lead to stress-induced apoptotic cell death. There are three major ER sensors that can initiate this system: PERK (Eif2 k3), ATF6, and IER1 (Ern1). IER1 was significantly increased in EtOH-0 mice compared to air controls and is associated with early time points of UPR activation, whereas ATF4 is considered a later phase of activation (Nishitoh, 2012) and was increased in EtOH-8h animals. ATF4 is a key activator of the transcription factor CHOP that was in the orange-yellow module), so we examined the level of ribosome associated CHOP mRNA and found that it was elevated in EtOH-8h compared to EtOH-0h in the sequencing data set. Using RTqPCR from RiboTag IP RNA in a new cohort of mice, CHOP RNA was increased after 24 hours of withdrawal (and nearly significantly after 8 hrs); this may represent a gradual increase as withdrawal progressed. CHOP translation may have important clinical implications as it is associated with the induction of apoptosis (Heilig et al., 2010), and UPR and CHOP have also been implicated in inflammatory processes associated with neurodegenerative disease models (Mhaille et al., 2008; Kamarehei et al., 2019), and after cocaine-induced microglial activation (Guo et al., 2015). The potential role of CHOP in microglia after ethanol exposure has received little attention, although it’s induction in peripheral tissues (Ji et al., 2005) and in brain (Badanich et al., 2011) has been well documented. In one case where ethanol increased CNS CHOP, it was found in neurons but not astroglia, although microglia were not studied (Badanich et al., 2011).

One limitation of the present study is that there are different temporal trajectories of transcriptome, translatome, and proteome over time (von Ziegler et al., 2022), and we selected specific times for investigation that may not have captured the full arc of UPR as withdrawal progressed. In the future, more detailed time courses of changes in microglial UPR will be important, as the time course of changes to microglial protein expression can occur over hours to days.

Based on the RNAseq results we examined the impact of CHOP cKO from microglia to determine its contribution to alcohol withdrawal and motivation to consume alcohol. We found that selective deletion of CHOP from microglia blunted several behavioral and physiological features associated with withdrawal after CIE. Altered thermoregulation, increased anxiety-like behavior, and increased alcohol consumption after CIE were all ameliorated by CHOP cKO, suggesting that CHOP activation in microglia during withdrawal has adverse consequences. This suggests that blocking CHOP signaling during withdrawal in individuals with an extensive history of alcohol consumption or current AUD may be therapeutic.

In summary, we identified alcohol-associated impairments in microglial function, including upregulation in the UPR cellular stress response and impaired metabolic function during early withdrawal. We found robust indicators of microglial activation, including an inflammatory cytokine response that evolves over the withdrawal timeline, as well as upregulated markers of active phagocytosis and apoptosis. By identifying affected gene networks and specific individual genes associated with acute intoxication and withdrawal in an alcohol dependence model, we have identified multiple novel points of possible therapeutic intervention. We focused on the UPR by testing CHOP cKO mice and found that deleting this transcription factor from microglia blunted some aspects of the complex response to alcohol withdrawal, suggesting that reduced activation of UPR in microglia may blunt the physiological, emotional, and relapse features associated with withdrawal from chronic alcohol exposure. Similarly, therapeutic modulation of microglia could possibly help attenuate the deleterious, proinflammatory, and cytokine-associated neurotoxicity characteristic of withdrawal. Ideally, either approach could be utilized to reduce the deleterious consequences of chronic alcohol use and withdrawal and thereby warrants further investigation.

## Methods

### Subjects

Experimental subjects for the RiboTag experiments were generated by crossing hemizygous Cx3cr1-Cre breeder mice (B6.129P2(C)-Cx3cr1tm2.1(Cre/ERT2)Jung/J, Stock #020940, The Jackson Laboratory, ME; Cx3cr1CreER) with homozygous Ribotag breeder mice (B6J.129(Cg)-Rpl22 tm1.1Psam/SjJ, Stock # 029977, The Jackson Laboratory, ME). Cx3cr1-Cre/Ribotag double transgenic mice (Figure 1a) were utilized for RNA-Seq (n=24) and qPCR (n=12) experiments to characterize the effect of ethanol intoxication and withdrawal on the striatal microglial translatome, with their Ribotag (Cx3cr1-Cre null) single transgenic littermates (n=14) utilized as genetic controls. To assess the influence of chronic alcohol exposure and subsequent withdrawal on protein and mRNA levels of key molecular markers of the UPR, male (n=36) and female (n=35) C57Bl/6J mice underwent CIE exposure and were sacrificed following their last ethanol exposure. A final group of 78 experimental subjects were generated by crossing hemizygous Cx3cr1CreER breeder mice with homozygous CHOPfl/fl breeder mice (B6.Cg-Ddit3tm1.1Irt/J, Stock #030816, The Jackson Laboratory, ME) to analyze the effect of CHOP conditional knockout in microglia on ethanol withdrawal-related behaviors. Twenty-two double transgenic Cx3cr1-Crewwt/tg/CHOPfl/fl mice were utilized for the Drinking in the Dark procedure, with 20 (Cx3cr1-Crewt/wt/CHOPfl/fl) single transgenic littermates (Cx3cr1-Crewt/wt/CHOPfl/fl) functioning as controls. Mice were group housed with same sex littermates (up to 5 per cage) and provided with ad-libitum food and water and a 14-10 light-dark cycle. All procedures were approved by the University of Washington Institutional Animal Care and Use Committee, and consistent with ethical guidelines in the Guide (National Research Council, 2011).

### Tamoxifen Induction of CRE

Four weeks before the start of the CIE paradigm, transgenic mice were injected with 75mg/kg Tamoxifen ip (20mg/mL tamoxifen in corn oil) once daily for five consecutive days at 14 weeks of age. Microglia specific Ribotag expression was confirmed in a subset of subjects using double label immunofluorescence for IBA-1 and HA (Figure 1b). Two additional Cx3cr1CreER-Ribotag mice were injected with only vehicle (corn oil) over 5 days, also underwent the CIE paradigm, and were used as Tamoxifen-negative controls for the RNA-Seq experiments.

### Chronic Intermittent Ethanol Paradigm

Starting at 18 weeks of age, subjects underwent 5 consecutive weeks of Ethanol Vapor or Air exposure following the CIE paradigm (Figure 1b) (Rose et al., 2016). Each week consisted of 4 CIE sessions where subjects were exposed to 16 hours of ethanol vapor (or air), from 6pm to 10am the following day. After every 4 CIE exposures, there was a 3-day break before starting the next week of sessions. 30 minutes before the start of each session, subjects were pretreated with an IP injection of either pyrazole alone (1mmol/kg bodyweight, in 0.9% saline) for Air treated subjects, or pyrazole (1mmol/kg) + ethanol (1.6g/kg; 10% ethanol in 0.9% saline) for Ethanol Vapor treated subjects. The ethanol vapor concentration in chambers was 12mg/L in order to consistently achieve sustained blood ethanol concentrations at approximately 200mg/dL in pyrazole/ethanol pretreated subjects (Figure S1). Food and water were available ad libitum during all sessions, including food at the bottom of the cage to enable feeding even when ataxic. Following every vapor exposure session, vapor exposed bedding and food was disposed and replaced with fresh bedding and food.

Two chambers were utilized, one for ethanol vapor treatment and the other for air control treatment, each with the dimensions of 32” W x 14” H x 21.5” D. Total air flow (40 SCFH) was matched between the two chambers during each session, and consistent across all sessions (2010). For the ethanol chamber, ambient air was bubbled through a side-arm flask containing 90% ethanol, producing ethanol vapors that were pumped into the chamber, and mixed with atmospheric air to achieve an ethanol vapor concentration of 12mg/L. Ethanol vapor concentration was measured using a breathalyzer (FC10; Lifeloc Technologies Inc.; Wheat Ridge, CO). Air pumps were turned on for 30 minutes before the start of each CIE session to confirm the ethanol vapor concentration. Excess air/vapor was ventilated into the rooms air exhaust intake.

### Necropsy and Tissue Processing

All mice utilized for RNA-Seq analysis were sacrificed by decapitation, brains were quickly dissected and then bisected into left and right hemispheres. For qPCR utilization, brains were quickly dissected and then bisected into left and right hemispheres immediately (n=3), 8 hours (n=5), or 24 hours (n=4) following their last ethanol exposure. Control animals (n=4) of similar age did not receive ethanol exposure. Whole striatum from the right hemisphere was microdissected, placed in 4°C supplemented homogenization buffer (50 mM Tris-HCl, 100 mM KCl, 12 mM MgCl2, 1% NP40, 1 mM DTT, 1× protease inhibitor cocktail (Sigma-Aldrich), 200 U/mL RNasin (Promega, Madison, WI), 100 µg/mL cyclohexamide (Sigma-Aldrich), 1 mg/mL heparin (APP Pharmaceuticals, Lake Zurich, IL)), and immediately homogenized. Homogenates were centrifuged at 10k RPM for 10 min, the supernatant collected and stored at -80°C until processed further for RNA extraction. For qPCR RNA extraction, the supernatant was collected with ten percent set aside as the whole transcriptome sample (input; IN) and the remaining sample set to be processed to isolate ribosome bound RNA (immunoprecipitate; IP). The left hemisphere was snap frozen and stored at -80°C.

### RiboTag RNA Isolation and Quantification

Striatal microdissections homogenized in the supplemented homogenization buffer were processed using our standard RiboTag procedure described in detail in our previous publication (Coffey et al., 2022). For qPCR analysis, RNA from both immunoprecipitated (IP) and input fractions were isolated using RNeasy Plus Micro Kit and eluted with 14–16 μl of water. RNA concentration was measured using Quant-iT RiboGreen RNA Assay (ThermoFisher Cat. R11490, Waltham, MA).

### RNA-Seq Library Preparation and qPCR

RNA-Seq libraries were prepared using SMARTer Stranded Total RNA-Seq Kit v2 – Pico Input Mammalian (Takara Bio USA, Inc. Cat. 635007, Mountain View, CA). 10 ng of IP RNA from each subject and 10 ng of pooled Input RNA from each treatment group was used to generate the libraries. RNA-Seq libraries were submitted to Northwest Genomics Center at University of Washington (Seattle, WA) where library quality control was measured using a BioAnalyzer, library concentrations were measured using Qubit dsDNA HS Assay Kit (ThermoFisher). Samples were normalized and pooled prior to cluster generation on HiSeq High Output for paired end reads. RNA-Seq libraries were sequenced on the HiSeq4000, Paired-end 75bp with PhiX spike-in controls (7%) (Illumina San Diego, CA). Negative control IP sequencing was also performed on an equivalent volume of eluate from two mice that did not receive tamoxifen, yeilding very low amounts of RNA.

RNA used in qPCR assays was reversed transcribed to create cDNA libraries for qPCR using Superscript VILO Master Mix (ThermoFisher Cat. 11,754,050, Waltham, MA). Following cDNA dilution, the qPCR assay was run using Power Sybr Green on the QuantStudio 5 Real-Time PCR System (ThermoFisher). Reactions were assembled for genes of interest (Ddit3, Akt2, Stip1). The normalized relative starting quantities (NRStQ) for gene expression was determined using the ΔΔCt method and normalized to three housekeeping genes (Ppia, Hprt, and Actb). Normalization factors were generated based on average housekeeping levels for IP and Input samples independently since RiboTag-IP RNA levels were consistently lower than Input. These values were analyzed using generalized linear mixed model. Primers for qPCR are as follows:

Ddit3 :

forward: 5’ CAGCGACAGAGCCAGAATAA

reverse: 5’ CACCGTCTCCAAGGTGAAA

akt2:

forward:5’CTGTCCTCTGGGCAATCTTATC reverse: 5’ CTCAGCTTCCAAAGGCTTCT

### Transcript Quantification and Quality Control

Raw fastq files were processed using multiple tools through the Galaxy platform (Afgan et al., 2016). Fastq files were inspected for quality using FastQC (Galaxy Version 0.7.0), and then passed to Salmon (Patro et al., 2017) (Galaxy Version 0.8.2) for quantification of transcripts. The Salmon index was built using the protein coding transcriptome GRCm38-mm10.

### Groups for Bioinformatics

Four groups were defined for differential expression analysis. Each group consisted of 6 mice (3 male, 3 female). Air 0h mice were exposed to air and sacrificed immediately after their final session. Air 8h mice were exposed to air and sacrificed 8 hours after their final session. Ethanol 0h mice were exposed to ethanol and sacrificed immediately after their final session. Ethanol 8h mice were exposed to ethanol and sacrificed 8 hours after their final session, while in withdrawal.

### Differential Expression Analysis

Differential gene expression was calculated using DESeq2 (Love et al., 2014) (Galaxy Version 2.11.39). All Salmon and DESeq2 settings were left default. To determine microglia specific gene enrichment, all IP samples were compared to all Input samples. For all other tests of differential expression, IP samples were compared to IP samples. To determine the effects of sacrifice time alone, Air 0h mice were compared to Air 8h mice; for the effects ethanol exposure, Ethanol 0h mice were compared to all Air mice; for the effects of withdrawal, Ethanol 8h mice were compared to Ethanol 0h mice. For all comparisons, a positive Wald statistic means that a gene is expressed more in the first group as compared to the second group. An FDR threshold of q = 0.05 was used throughout the manuscript.

### Gene Set Enrichment Analysis

Wald statics generated by DeSeq2 were used as the ranking variable for gene set enrichment analysis. All genes with reliable statistical comparisons (those not filtered by DeSeq2) were entered into WebGestalt 2019 (Liao et al., 2019) and GSEA was run all pertinent comparisons (enrichment, IP Air, IP Ethanol, IP Withdrawal). Gene sets analyzed include GO: Molecular Function, Reactome, Wiki Pathways, and Transcription Factor Targets. All advanced parameters were left default except for significance level, which was set to FDR = 0.05.

### Weighted Gene Co-Expression Network Analysis

A topological overlap matrix for IP samples was generated, and module clustering was accomplished using the WGCNA (Langfelder and Horvath, 2008) package for R (R Development Core Team, 2010). Briefly, the TPM matrix for each group was filtered to remove zero-variance genes, and a signed adjacency matrix was generated using “bicor” as the correlation function. From this a signed topological overlap matrix was generated, followed by a dissimilarity topological overlap matrix. Finally, module membership was assigned using a dynamic tree cut. Clusters produced from this procedure are referred to as gene modules and randomly renamed using the “Crayola Color Palette” in order to combat attribution of meaning or importance to numbered modules (Cobeldick, 2020). Modules were merged by clustering on module eigengenes with a cut height of 0.85. A custom WGCNA MATLAB Class was used for graphing gene networks (GitHub.com/DrCoffey/WGCNA).

### Fluorescent in-situ Hybridization for CHOP and DNAJA1

Adult male (n=15) and female (n=15) C57Bl/6J were deeply anesthetized with an intraperitoneal injection of sodium pentobarbital and phenytoin sodium and perfused transcardially with 0.1 M phosphate-buffered saline (PBS), followed by cold 4% paraformaldehyde (pH 7.4) immediately or 8 hours following their last ethanol (or air) exposure. Brains were dissected, post fixed in 4% paraformaldehyde overnight at 4°C and transferred to 30% sucrose in PBS. Sections (16µm) across the rostro-caudal axis of the striatum were collected on a Leica CM3050 S cryostat, mounted on charged slides, and dehydrated at -20°C for 30-minutes before being frozen at -80°C. The RNAscope (ACD Bio) fluorescent multiplex in-situ hybridization was used to visualize and quantify Ddit3 (CHOP) and Dnaja1 (DNAJA1: heat shock protein chaperone) mRNA in the striatum for each animal. Slides were set up so that an animal from each group would have similar sections for each slide to reduce inter-slide variability. Following RNAscope protocol, to localize mRNA to microglia, immunohistochemistry was performed using primary anti-Iba1 rabbit antibody diluted 1:1000 (FUJIFILM Wako; #019-19741) overnight at 4ºC and secondary Alexa Fluor 488 goat anti-rabbit diluted 1:400 (ThermoFisher; #A-11008). Sections were then washed in PBS (3x, 10 minutes) and cover slipped with Pro-Long Gold with DAPI mounting medium (ThermoFisher; #P36931). Negative controls had only probe diluent applied and were stained via IHC to further characterize morphology of microglia without the RNAscope probes.

### Immunohistochemistry for CHOP, grp78, IRE1α, Akt2, and Stip1

Adult male (n=25) and female (n=24) C57Bl/6J were deeply anesthetized with an intraperitoneal injection of sodium pentobarbital and phenytoin sodium and perfused transcardially with 0.1 M phosphate-buffered saline (PBS), followed by cold 4% paraformaldehyde (pH 7.4) immediately or 8 hours following their last ethanol (or air) exposure. Brains were dissected, post fixed in 4% paraformaldehyde overnight at 4°C and transferred to 30% sucrose in PBS. Sections (18µm) across the rostro-caudal axis of the striatum were collected on a Leica CM3050 S cryostat, mounted on charged slides, and dried at -20°C for 30-minutes before being frozen at -80°C. On-slides sections were thawed, rinsed in 1X PBS (3X, 10 minutes), permeabilized with 0.5% Triton PBS and then underwent heat-induced epitope retrieval in citrate buffer at 99°C for 10 minutes (ACD Bio; #322000), and blocked with 4% BSA in 0.3% Triton PBS at room temperature for 1 hour.

For immunohistochemistry of Iba1 and either CHOP, grp78, or IRE1α, on-slide sections were incubated with primary anti-Iba1 mouse antibody diluted 1:2000 (Abcam; #ab283319) with either anti-CHOP rabbit antibody diluted 1:250 (Sigma Millipore; #SAB5700602), anti-grp78 rabbit antibody diluted 1:250 (Sigma Millipore; #SAB4501452), or anti-IRE1α rabbit antibody diluted 1:250 (Sigma Millipore; #SAB5700519) in 4% BSA and 0.3% Triton PBS for 32h at 4°C. For immunohistochemistry of Iba1, Akt2, and Stip1, on-slide sections underwent the same pretreatment before addition of primary anti-Iba1 rabbit antibody diluted 1:1000 (FUJIFILM Wako; #019-19741) with either anti-Akt2 mouse antibody diluted 1:250 (Santa Cruz Biotechnologies; #sc-5270) or anti-Stip1 mouse antibody (Santa Cruz Biotechnologies; # sc-390203) in 4% BSA and 0.3% Triton PBS for 24h at 4°C. Sections were then rinsed with PBS (3X, 10 minutes), incubated with both a 1:400 dilution of Alexa Fluor 488 Goat anti-rabbit secondary antibody (ThermoFisher; #A-11008) and 1:400 dilution of Alexa Fluor 567 Goat anti-mouse (ThermoFisher; #A-11004) for sixty minutes at room temperature, washed with PBS (3X, 10 minutes) and cover slipped with Pro-Long Gold with DAPI mounting medium (ThermoFisher; #P36931). For negative controls, primary antibodies other than Iba1 antibodies were omitted from the incubation.

### Image Processing and Analysis

Confocal image z-stacks were acquired for both in-situ and IHC sections in separate sessions on a Leica SP8X confocal microscope (Leica Microsystems) at 40x magnification. Using Imaris (Bitplane), the images were processed by applying background and Gaussian filtering to all channels. Three-dimensional ‘surfaces’ of the microglia were then created by filtering out any signal in the color channel associated with microglia with a volume larger than 0.01µm3. Three-dimensional ‘spots’ were created for each mRNA fluorescent signal for both CHOP and DNAJA1 molecules. Spot-to-surface colocalization (RNAscope) and intensity-based colocalization (RNAscope and IHC) were used to quantify the overlap of targets with microglia. A Sholl Analysis plugin along with the AI-assisted filament creator was used to assess microglia morphology (e.g. Sholl interactions, filament branching, process end points) in each of the sections. Separate intensity-based colocalization was done through custom MATLAB app (https://github.com/DrCoffey/fishFinder) to verify the results seen using Imaris. Statistical analysis of the data generated was done with Graphpad Prism and JMP. Initial 3 way generalized linear model revealed no sex differences; subsequent analysis was performed with a 2×2 design comparing vapor treatment and withdrawal time.

### Physiological and Behavioral Experiments

Cx3cr1-Cre/CHOP double transgenic mice (n=11) were utilized to analyze the effect of CHOP conditional knockout in microglia on ethanol withdrawal-related behaviors following CIE exposure, with CHOP (Cx3cr1-Cre WT) single transgenic (n=11) and hemizygous littermates (n=14) functioning as controls. Weights were taken three times a week to ensure subject health. To measure the effects of CIE exposure on thermoregulation, core body temperatures of experimental mice were measured using a rectal probe connected to a digital thermometer (World Precision Instruments; #RET-3 and #BAT-12R). The probe was inserted 2cm into the colon for a period of no longer than 30 seconds. Mice were habituated to temperature measurements before starting CIE to mitigate handling stress. Temperature measurements were then taken before the first alcohol exposure of the week (baseline), immediately following the last exposure of each week (wAM), and 8 hours later during the height of withdrawal (wPM), for the duration of the CIE exposure.

#### Sucrose Preference

Animals were exposed to sucrose and water bottles in their home cages for three days prior to this session to habituate to the bottles and sucrose solution. To assess hedonic sensitivity, mice were placed in 2-bottle choice lickometer chambers (47cm x 25cm x 20cm) containing a bottle of 2% sucrose solution or water for 3 hours as a baseline measurement before being administered tamoxifen or CIE exposure. All animals demonstrated sucrose preference of greater than 75% in their baseline sessions prior to being exposed to ethanol and subsequent experiments. The test session lasted 3 hours, the placement of the sucrose bottle on either the right or left side of the chamber was counterbalanced between animals, and the licks were automatically counted. The metrics included the total number of licks for either sucrose or water and the percentage of licks on the sucrose bottle compared to total licks of both bottles.

#### Open Field

To analyze the effects of CIE exposure on anxiety-like phenotypes, 48 hours after the last exposure to ethanol (or air), mice were allowed to freely explore an open field chamber (40cm x 32cm x 24cm) while being video recorded for 30 minutes. Locomotion, percentage of time spent in the center area, latency to approach the center, and immobility were all scored and analyzed using Ethovision (Noldus Technologies).

### Drinking in the Dark (DID)

A new cohort of thirty-nine C57Bl/6 microglia CHOP conditional knockout (cKO) (n=20) and littermate control (n=19) mice began the Drinking in the Dark (DID) procedure. The overall experimental structure included an ethanol consumption pretest, CIE vapor exposure, and ethanol consumption posttest (Figure 5a). Before beginning DID, the mice gradually acclimated to a new light cycle by shifting the cycle forward by 2 hours per week for 8 weeks, stopping when the light cycle was shifted from the previous 6 A.M. to 8 P.M. 14 h light cycle, to the new 10 P.M. to 12 Noon 14 h light cycle. Four weeks before the start of the CIE paradigm, Cx3cr1-Cre/CHOP mice were injected I.P. with 75mg/kg Tamoxifen (20mg/mL tamoxifen in corn oil) once daily for five consecutive days to induce CRE expression.

#### Ethanol Consumption Testing

We used a modification of the DID procedure (Thiele and Navarro, 2014). After adjusting to the new light cycle for an additional week, the mice began a baseline ethanol drinking pretest. Three hours into the dark cycle, the mice were removed from their home cages and placed into one side of a new cage; each cage was split with a cage divider, with two mice per cage. For each side of each cage, a single Bio-Serv liquid feeding tube (Bio-Serv, product #9019, Flemington, NJ) containing 20% ethanol in Hydropac™ water matching the water in the home cages was weighed and placed inside. With all the lights off, the mice were left for 2 hours, and at the end of this period consumption was measured by reweighing each bottle and subtracting it from the initial weight. This pretest procedure was repeated for 3 consecutive days.

Following the 3 days of pretest and 2 days of rest, the mice began 5 weeks of CIE exposure in the ethanol vapor chamber as described above. Mice were preassigned to ethanol vapor (microglia CHOP cKO (n=9), controls (n=9)) or air (microglia CHOP cKO (n=11), controls (n=10)) conditions in advance of the ethanol consumption pretest. In this cohort we gradually escalated the pre-vapor ethanol and pyrazole injections (Figure 7a) which allowed for a gradual increase of ethanol tolerance over the 5 weeks of vapor exposure, with a maximum dose of 0.5 ml of 10% ethanol with 3.45 mg/ml pyrazole, in sterile saline. Three nights after the completion of week 5 of the CIE exposure, the mice were once again placed in cages with bottles of 20% ethanol, repeating the same 2h ethanol consumption procedure used for the baseline pretest. Following the final consumption test day, the mice were transcardially perfused with ice-cold phosphate-buffered saline (PBS) followed by 4% paraformaldehyde (PFA) for potential future analysis. Brains were post-fixed at 4°C for 24 h in 4% PFA before being transferred to 30% sucrose in PBS. The mean consumption over the three days for each test session was used for analysis; cages that had visible liquid spills in the cage at the end of a session were omitted on that day, and the remaining days for that cage were used to determine mean consumption expressed as grams ethanol per kilogram body weight (g/kg). Mean consumption data was analyzed by generalized linear mixed model analysis with pre and post vapor exposure as repeated measures, with genotype (microglia CHOP cKO vs control) and exposure (ethanol vapor vs air) as between-subjects factors.

### Experimental Design and Statistical Analysis

The overall experimental design is illustrated in Figure 1. Statistical analysis for behavior and image analysis was performed using either Prism 10 or JMP Pro 18 statistics package. Statistical analysis was performed using generalized linear models (GLM) or generalized linear mixed models (GLMM) with sex, genotype, and treatment condition being the between-group factors. Where appropriate, Tukey’s post-hoc test was applied between treatment groups and Dunnet’s post-hoc test was applied when there was a defined control group over time. Differential gene expression analysis was performed using DESeq2. Gene sets were analyzed using WEB-based GEne SeT AnaLysis Toolkit (www.webgestalt.org). All data and code used to generate the figures in this manuscript is available in our GitHubRepository. Raw sequencing and image data prohibitively large to host so it is available upon reasonable request.

## Acknowledgments

Supported by R21 DA044757, T32 DA007278, and T32 AA007455.

## Author Contributions

Rapheal Williams, Brett Dufour, William Nickelson, Atom Lesiak, Aliyah Dawkins performed the experiments, analyzed the data, and wrote the manuscript. Kevin Coffey analyzed the data and wrote the manuscript. Atom Lesiak performed the experiment and analyzed the data. Gwenn Garden provided animals, assisted in data interpretation, and wrote the manuscript. John Neumaier designed the experiment and wrote the manuscript.

## Current Affiliations

Rapheal Williams: Gerontology, Biological Sciences, Biochemistry and Molecular Medicine, University of Southern California, Los Angeles CA 90089, USA; Atom Lesiak: Department of Genome Sciences3, University of Washington School of Medicine, Seattle, WA, 98104, USA; Gwenn A. Garden: Department of Neurology4, University of North Carolina Chapel Hill, NC, 27599, USA; Brett Dufour: Department of Pathology and Laboratory Medicine5, University of California Davis, CA, 95616, USA; Aliyah J. Dawkins: Department of Neurobiology, Harvard University, Boston, MA 02115, USA; William B. Nickelson: Neuroscience Graduate Program, University of Southern California, Los Angeles CA 90089, USA.

---

**Figure S1.**
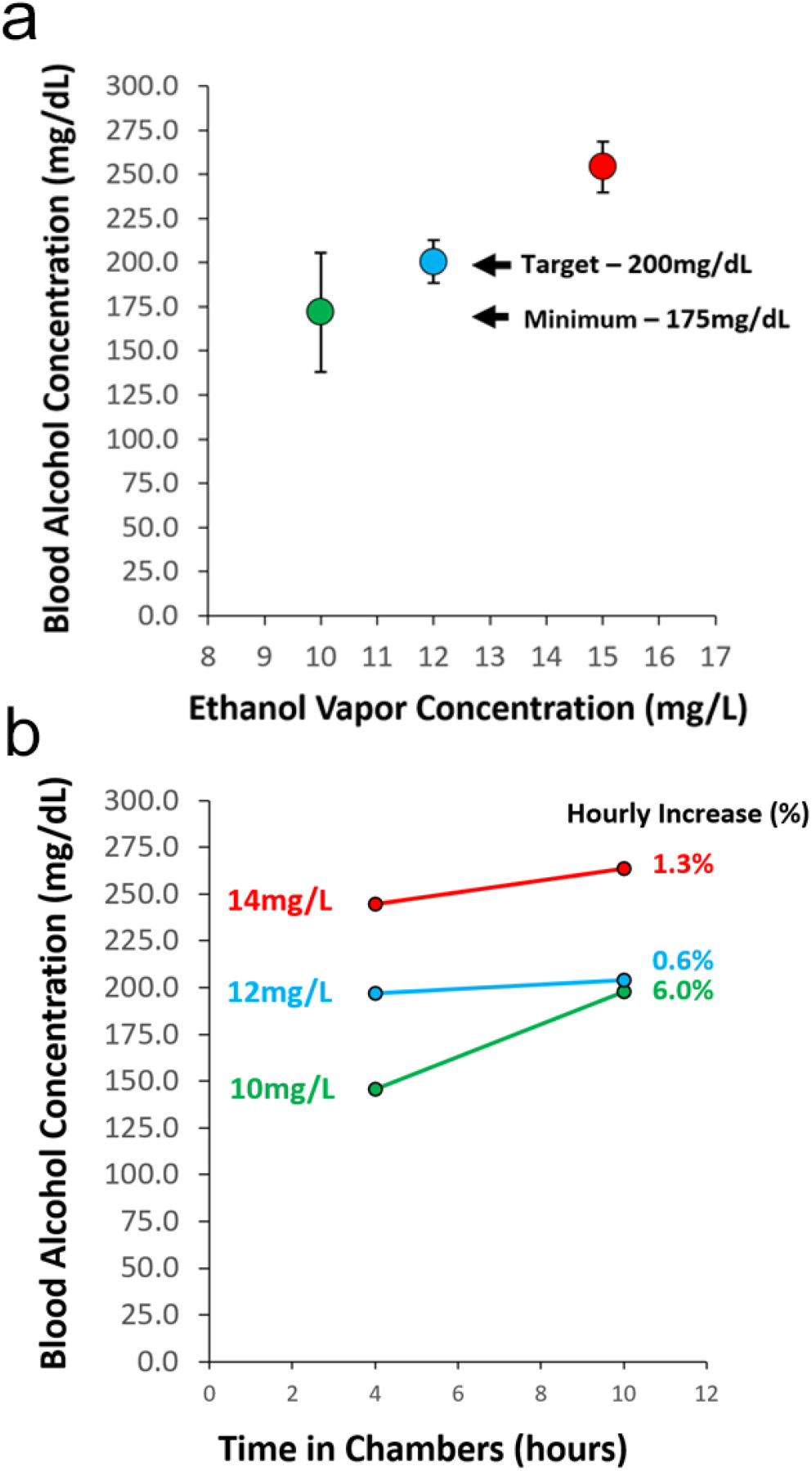
Blood Alcohol Stability. **a)** Blood alcohol concentration is closely related to ethanol vapor concentration allowing us to tightly control blood alcohol concentration around 200 mg/dL. **b)** At 12 mg/L ethanol vapor concentration, blood alcohol concentration remains steady across all 10 hours in the chamber.

**Figure S2.**
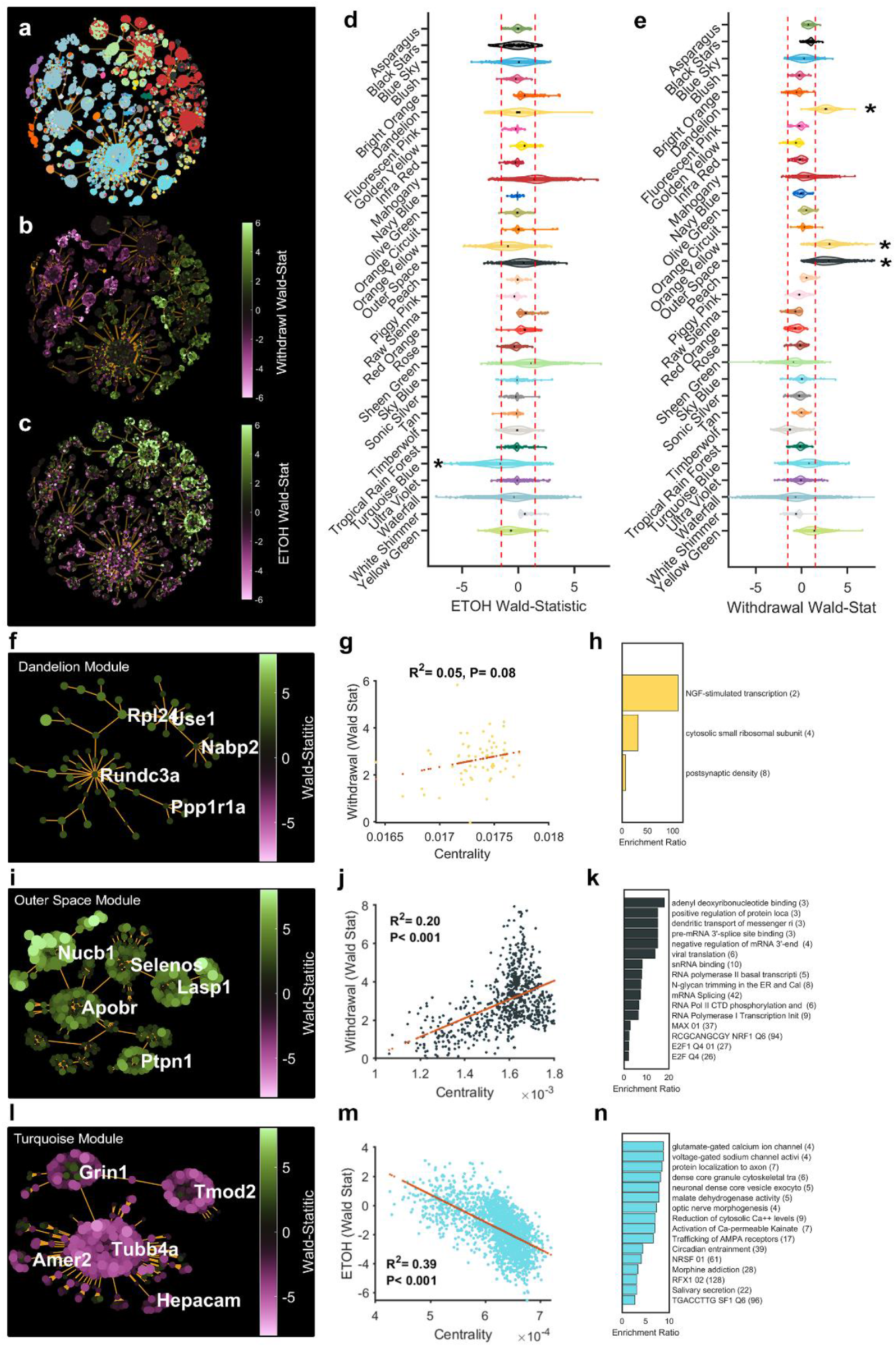
Complete WGCNA Analysis. **a)** Genes are coded by module color and projected into a 2D minimum spanning tree using a “force” layout. **b)** The same genes are coded by differential expression (Wald-statistic) data during ethanol exposure and **c)** ethanol withdrawal. WGCNA modules are plotted based on their differential expression metrics (Wald-statistic) for **d)** ethanol exposure and **e)** ethanol withdrawal. Modules who’s mean Wald-statistic was less than -1.5 or greater than 1.5 were flagged for further analysis. **f)** The Dandelion module projected into 2D space using a minimum spanning tree and showing the top five most central genes. **g)** Centrality (a measure of network importance) in the Dandelion module is not correlated to differential expression for withdrawal. **h)** Over-representation analysis on the Dandelion module shows that genes related to NGF signaling were upregulated after ethanol withdrawal in striatal microglia. **i)** The Outer Space module projected into 2D space using a minimum spanning tree and showing the top five most central genes. **j)** Centrality in the Outer Space module is correlated to differential expression for withdrawal. **k)** Over-representation analysis on the Outer Space module shows that genes related to RNA splicing, transcription, and polymerization were upregulated after ethanol withdrawal in striatal microglia. **l)** The Turquoise module projected into 2D space using a minimum spanning tree and showing the top five most central genes. **m)** Centrality in the Turquoise module is negatively correlated to differential expression after ethanol exposure. **n)** Over-representation analysis on the Turquoise module shows that genes classically associated with neurons were downregulated in striatal microglia after ethanol exposure.

**Figure S3.**
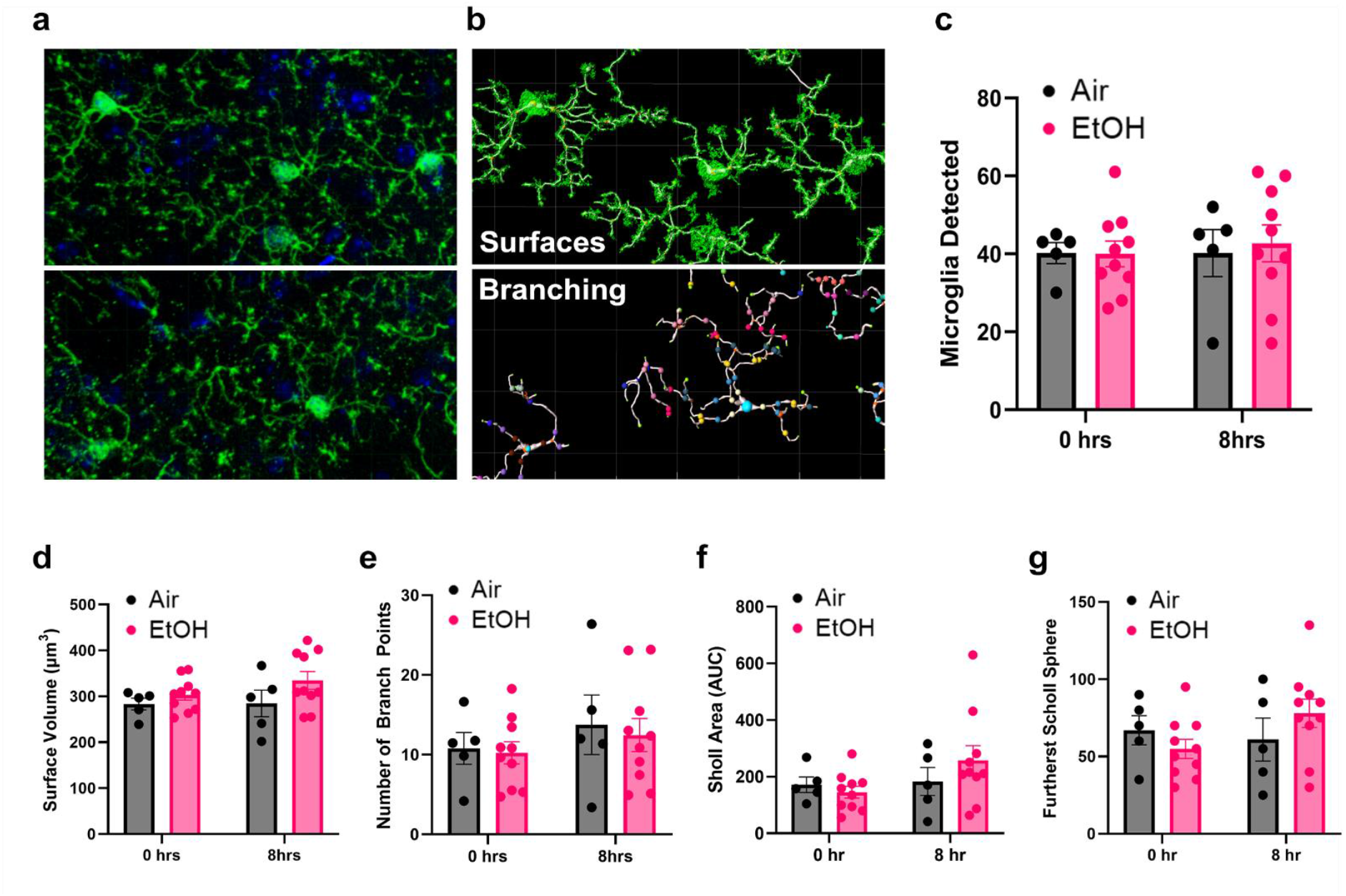
CIE effects on microglia morphology. Within the striatum using Iba1 staining. This was **a)** followed by construction of 3D filaments using the masked microglial structures of each image as a base **b)**. There were no significant differences in the number of microglia **c)** and their surface volumes **d)**, microglia branch points **e)**, and Sholl interactions between withdrawal time points and treatment groups **f**,**g)**. All columns represent mean values calculated for each subject animal from several tissue sections analyzed for each animal.

**Figure S4.**
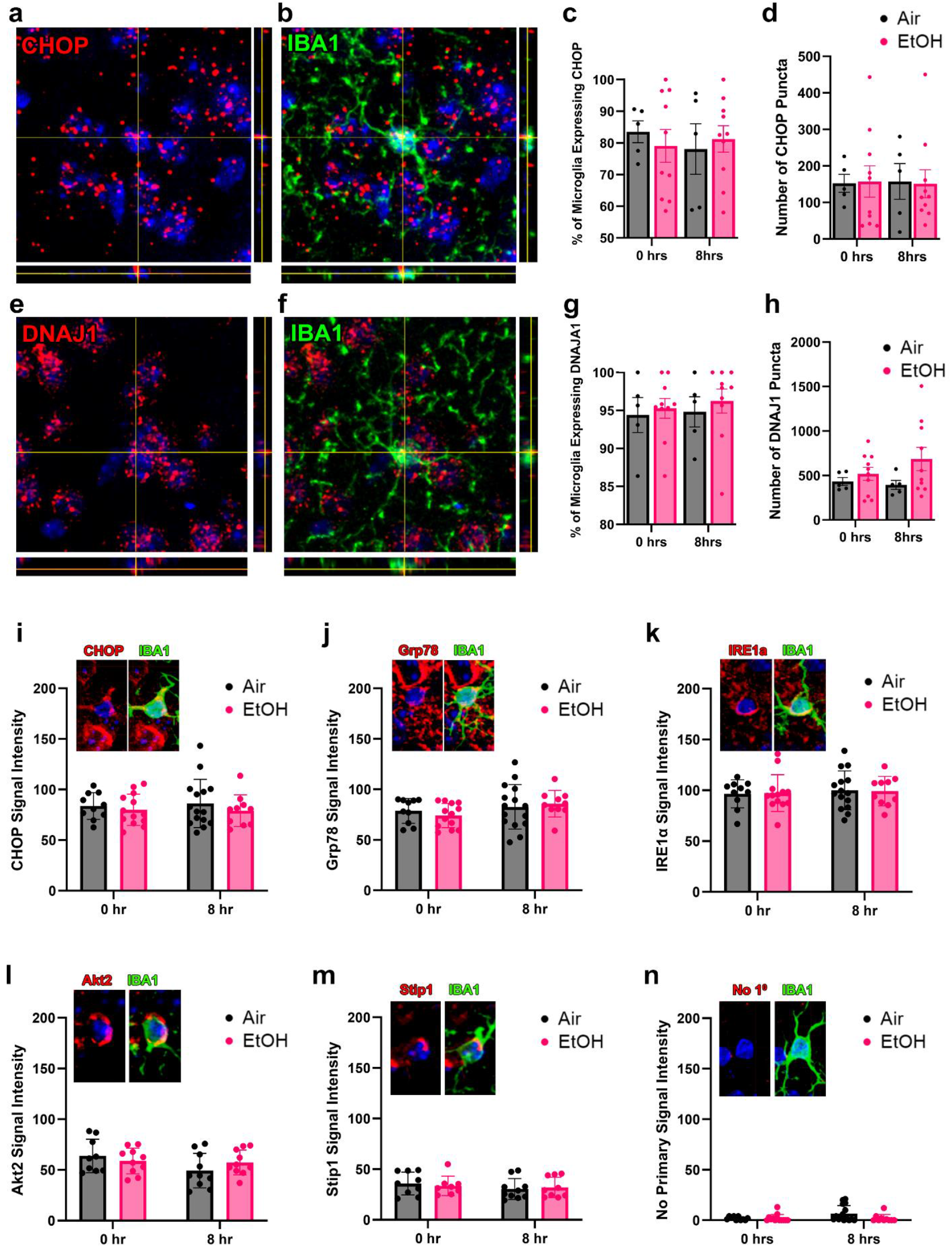
In-situ Hybridization and immunohistochemistry for CHOP and Other UPR Candidates. **a, b)** CHOP (Ddit3) and a top enriched gene Dnaja1 **e, f)** were measured using RNAscope performed in combination with Iba1 immunohistochemistry to label microglia **b, f)**. The percent of microglia expressing CHOP **c)** and, Dnaja1 **g)** did not change during the withdrawal timepoint. The number of RNA-associated puncta were counted within each microglia for CHOP **d)** and Dnaja1 **h)**. Immunohistochemistry was performed and pixel intensity within microglia was measured using custom intensity-based colocalization MATLAB scripts. Pixel intensities for CHOP **i)**, Grp78 **j)**, IRE1α **k)**, Akt2 **l)** and Stip1 **m)** within Iba1-labeled microglia did not show differences between withdrawal timepoints for either air or ethanol groups, suggesting that any changes in these protein levels do not develop in the first 8 hours of withdrawal. Separate channels for Iba1 and the proteins of interest overlapped with DAPI are separated and merged for ease of viewing in within-graph insets. All columns represent mean values calculated for each subject animal from several tissue sections analyzed for each animal.

